## Supplemental Tables and Figures in one pdf file for "Stem Cell Transplantation Rescued A Primary Open-Angle Glaucoma Mouse Model"

### Supplementary Figures

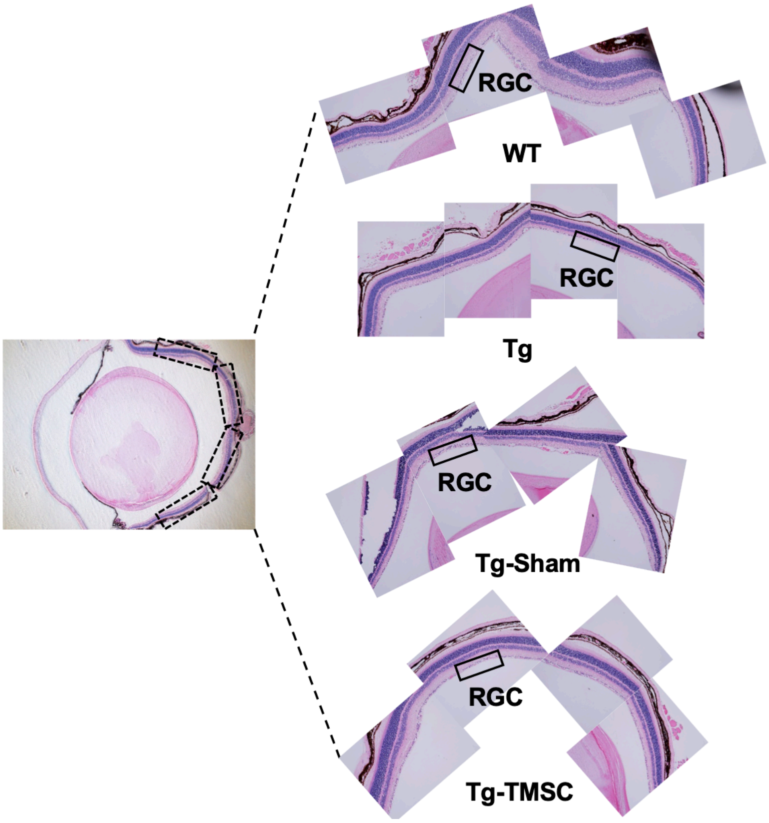


**Figure S1: Transplanted TMSCs rescue RGCs and prevent neurodegeneration in Tg-MyocY437H mice.** Eye sections stained hematoxylin and eosin show the RGC layer in the eyes. The black INSET boxes in the left picture show the areas from which RGCs were captured so that the RGCs in the whole retina were counted in the central sections containing optic nerve.

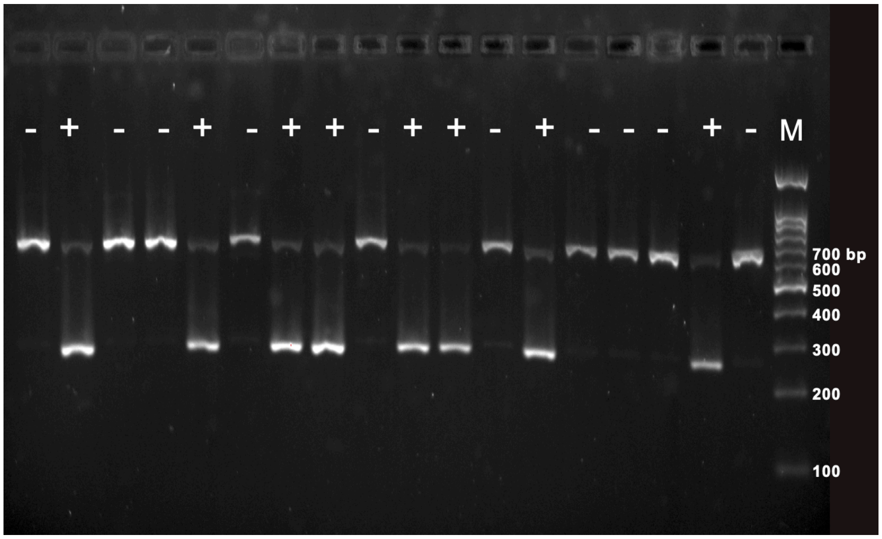


**Figure S2. Genotyping of transgenic mice by polymerase chain reaction (PCR).** Tg-Myoc Y437H mice (+) displayed with PCR products of 249 bp (MyocY437H) and weak band at 610 bp (mouse DNA). The mice with only 610 bp band were transgenic negative mice (-). M: DNA markers.

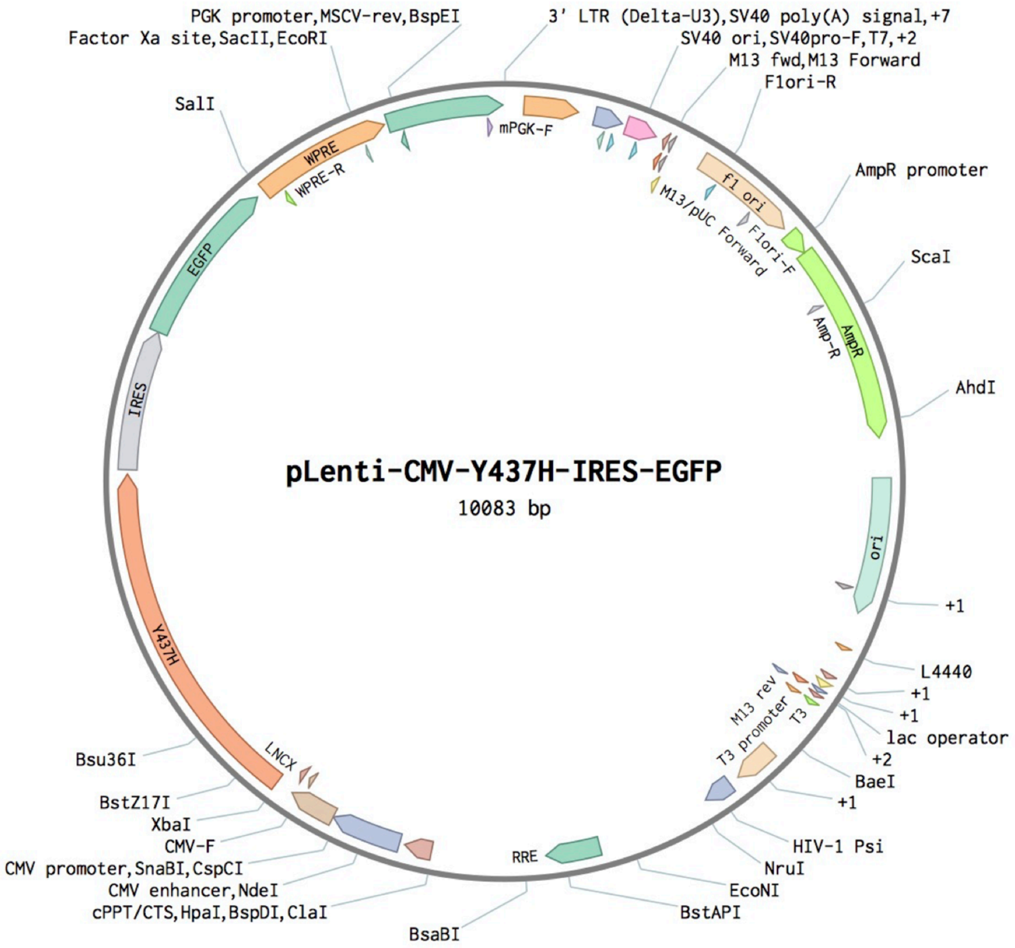


**Figure S3. Structure of lentiviral packaging plasmid: pLenti$^{CMV\text{-}Y437H\text{-}IRES\text{-}GFP}$**

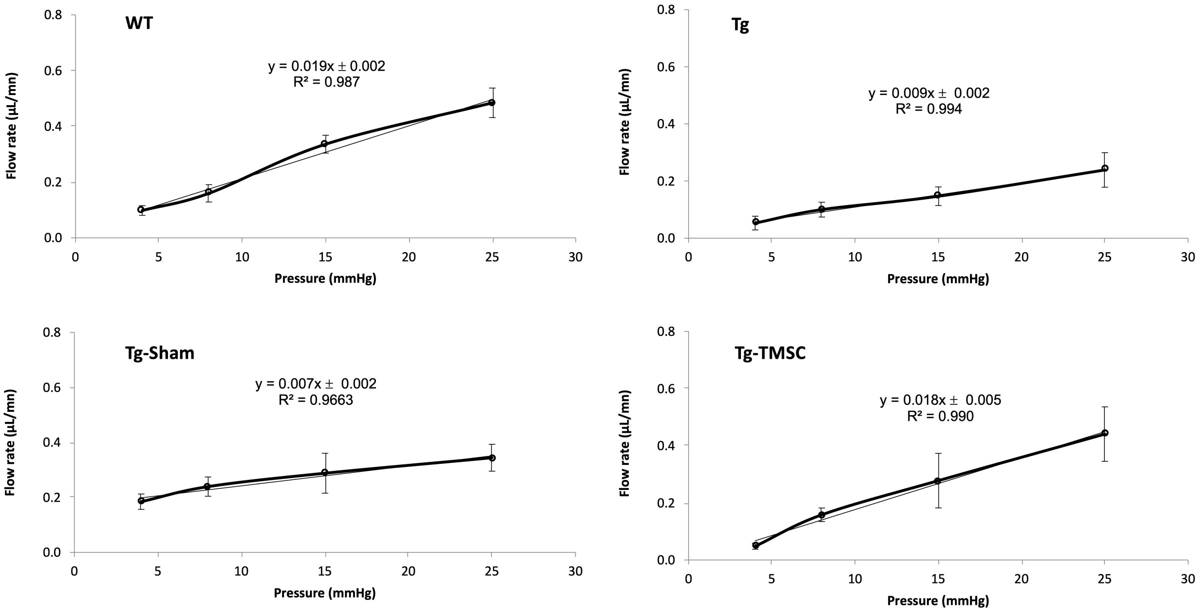


**Figure S4. Representative perfusion outflow data from each mouse group.** X-axis: perfusion pressure (mmHg), Y-axis: flow rate (µL/min), y is the slope indicating outflow facility (µL/min/mmHg).

**Supplementary Table 1.** Related gene expression increase ($p<0.05$) in genes related to TM ECM maintenance, TM integrity and motility in three individual TMSCs as compared to fibroblasts. Related to Figure 9A.

| | **Fibro-3** | **Fibro-2** | **Fibro-1** | **TMSC-3** | **TMSC-2** | **TMSC-1** |
|---|---|---|---|---|---|---|
| **ITGA3** | 4260.60416 | 5259.61544 | 4221.22963 | 19514.1613 | 5667.24335 | 10743.1066 |
| **CHI3L1** | 48.40425 | 351.5922 | 22.12826 | 2131.658 | 482.7891 | 1541.959 |
| **VTN** | 39.13535 | 26.4968 | 52.68634 | 36.58226 | 207.3045 | 654.0196 |
| **LOX** | 8841.50028 | 10320.5049 | 12332.8187 | 635.740272 | 564.789494 | 3450.54988 |
| **FST** | 5435.695 | 2775.031 | 5324.482 | 47082.35 | 9800.434 | 8603.46 |
| **COL4A6** | 189.4975 | 66.24201 | 80.08324 | 4491.708 | 5919.694 | 1349.094 |

**Supplementary Table 2.** Related gene expression increase ($p<0.05$) in genes related to TM ECM interaction.

| | **Fibro-3** | **Fibro-2** | **Fibro-1** | **TMSC-3** | **TMSC-2** | **TMSC-1** |
|---|---|---|---|---|---|---|
| **VTN** | 39.13535 | 26.4968 | 52.68634 | 36.58226 | 207.3045 | 654.0196 |
| **COL4A5** | 1822.884 | 852.9932 | 1141.186 | 7856.286 | 8172.403 | 3951.806 |
| **MYLK** | 2636.487 | 2302.165 | 1633.277 | 1357.498 | 397.1032 | 4418.69 |
| **PDGFD** | 2599.411 | 868.2799 | 2079.003 | 125.5661 | 3435.726 | 5045.021 |
| **COL4A6** | 189.4975 | 66.24201 | 80.08324 | 4491.708 | 5919.694 | 1349.094 |
| **HSPG2** | 11474.9 | 9403.308 | 17417.05 | 6801.333 | 8018.537 | 17596.47 |
